# Rapid and Interpretable AMR Diagnostics via Genomics and Cell Painting using Differential Geometry-based Directed-Simplicial Neural Networks on Multimodal Data

**DOI:** 10.64898/2026.03.11.711128

**Authors:** Lokendra S. Thakur, Sonia Shinde Mahajan, Gurpreet Bharj, Mengyuan Ding, Nino Dekanoidze, Vidhiti Shrivastava

## Abstract

Antimicrobial resistance (AMR) remains a critical global health challenge, particularly in high-prevalence regions such as India, where rapid and interpretable diagnostic tools are urgently needed. To address this challenge, we present a computational framework for AMR prediction that integrates genomic and cellular phenotypic data using an in-house developed differential geometry–based Directed Simplicial Neural Network (Dg-Dir-SNNs) applied to multimodal datasets.

Using this framework, we analyzed 384 clinically relevant AMR isolates, including *Escherichia coli* and *Klebsiella pneumoniae*, integrating 256 genomic k-mer features with 503 cellular morphology descriptors derived from high-content Cell Painting assays. The Dg-Dir-SNNs model constructs an inferred-causal network of top-ranked biomarker-driving features, predicting potential directional dependencies among genomic motifs and phenotypic features. Network analysis identified `kmer_TATG` as the top-ranked driver associated with predicted resistance, with a local neighborhood including other genomic motifs (`kmer_TTTT`, `kmer_CGTG`, `kmer_TCAC`, `kmer_CGTA`, `kmer_GAAA`, `kmer_TAAA`, `kmer_TACA`, `kmer_TGTG`, `kmer_TGAG`, `kmer_AAAA`) and a key morphological feature (`Cells_correlation_ER_Brightfield`). These relationships suggest potential mechanistic associations in which specific genomic motifs may influence cellular phenotypes linked to antimicrobial resistance.

Although not yet clinically deployed, this approach demonstrates the potential of multimodal AI-driven modeling for rapid *in silico* AMR prediction. By providing interpretable, biologically grounded insights, the framework may support future diagnostic development, targeted surveillance strategies, and experimental validation in high-resistance healthcare settings.

## 1 Background and Rationale

Antimicrobial resistance (AMR) constitutes a major global health crisis, threatening public health, economic stability, and the efficacy of modern medical interventions. In 2019, AMR was directly responsible for 1.27 million deaths worldwide, with an additional 4.95 million deaths associated with drug-resistant infections,^**1,2**^ highlighting its immense burden on healthcare systems. Factors such as antibiotic overuse, inadequate infection control, and poor sanitation accelerate resistance, compromising treatments for common infections and complex procedures including surgeries and cancer therapy, while inflating healthcare costs.^**2, 3**^ In India, pathogens like *Escherichia coli* (*E. coli*) and *Klebsiella pneumoniae* (*K. pneumoniae*) exhibit alarming multidrug resistance, complicating management of urinary tract infections, bloodstream infections, and pneumonia.^**4**^ Traditional culture-based diagnostics, though accurate, require 24–72 hours, delaying critical therapeutic decisions.^**1, 5**^ To address these limitations, we developed the **Differential Geometry–based Directed Simplicial Neural Networks (Dg-Dir-SNNs) pipeline** (see Figure 2), a unified geometric deep learning framework that integrates intrinsic manifold 1 learning, topology-aware feature refinement, and higher-order neural message passing. By jointly optimizing predictive performance, numerical stability, and interpretability, Dg-Dir-SNNs captures complex interactions among genomic, phenotypic, morphological imaging (cell-painting), and immune features while maintaining a consistent internal state (neighborhood graph **G** and simplicial complex tri) for reliable predictions. Beyond accelerating decision-making compared to traditional diagnostics, the model employs graph-based imputation and redundancy compression to handle missing or noisy data, and supports interpretability through SHAP and inferred-causal analyses. By enabling precise, actionable predictions for AMR, Dg-Dir-SNNs has the potential to significantly improve patient outcomes, guide targeted antimicrobial therapy, and inform public health strategies, thereby addressing both clinical and societal challenges posed by the global AMR crisis.

## 2 Study Design and Objectives

Antimicrobial resistance (AMR) is one of the most urgent and rapidly growing threats to global public health. The increasing prevalence of resistant pathogens, including *Escherichia coli* (*E. coli*) and *Klebsiella pneumoniae* (*K. pneumoniae*), particularly in high-burden regions such as India,^**4**^ complicates the treatment of common infections such as urinary tract infections, bloodstream infections, and pneumonia. These pathogens exhibit alarming levels of multidrug resistance, resulting in treatment failure, prolonged hospitalization, increased mortality, and escalating healthcare costs.

Current AMR diagnostics primarily rely on culture-based methods requiring 24–72 hours to produce results. Such delays hinder timely therapeutic decision-making and contribute to the continued spread of resistant strains. Addressing these limitations requires computational frameworks capable of rapidly predicting AMR patterns while providing clinically interpretable insights.

To address this need, this study develops a data-driven predictive framework based on Differential Geometry–Directed Simplicial Neural Networks (Dg-Dir-SNNs). The approach is designed to integrate multi-modal biological datasets, including genomic, phenotypic, immunological, and high-content cell painting assay data, to construct a unified predictive system for antimicrobial resistance. Directed Simplicial Neural Networks (Dir-SNNs),^**6**^ grounded in differential geometry, enable modeling of higher-order relationships across heterogeneous modalities and facilitate real-time prediction of resistance mechanisms.

The study is structured around two primary objectives. First, a comprehensive multi-modal framework is constructed by integrating genomic, phenotypic, immune profiling, and morphological imaging datasets into a scalable and high-quality analytical resource. Representative data sources (see Table. 1) include genomic repositories, phenotypic resistance databases, immune profiling datasets, and high-content imaging collections. Dataset quality is evaluated in terms of accuracy, completeness, and scalability, and predictive performance is compared against baseline machine learning approaches such as Random Forest, Logistic Regression, and conventional Neural Networks. The Dg-Dir-SNNs architecture is trained, assessed, and optimized through comparative analysis, with validation conducted using pathogen-specific case studies involving *E. coli* and *K. pneumoniae*. The model has been implemented and tested on integrated real datasets.

**Table 1.** Data sources, characteristics, and sample types relevant for AMR diagnostics, therapeutics, and predictive modeling, including public cell painting profile repositories.

| Data Type | Repository/Source | Description | Key Characteristics for AMR Predictions | Sample Type |
| --- | --- | --- | --- | --- |
| Genomic Data | NCBI (SRA) | Genetic sequences of AMR-associated genes in *E. coli* and *K. pneumoniae*. | Genome completeness, sequencing coverage/depth, annotated AMR genes, strain metadata, resistance mechanism classification (e.g., β-lactamases, efflux pumps). | Isolated DNA from bacterial cultures / clinical isolates |
| Phenotypic Data | PATRIC | Antibiotic resistance profiles, including MIC values for AMR pathogens. | Number of antibiotics tested, strain diversity, MIC measurement methods/units, geographic/temporal metadata, direct resistance phenotype labels. | Bacterial cultures, clinical isolates |
| Cell Painting Data | Broad Institute Cell Painting Gallery | High-content imaging capturing cellular morphology under drug exposure and genetic perturbations. | Cell morphological features extracted under perturbation, imaging resolution, staining channels/dyes, drug/perturbation conditions, morphological feature extraction. | Mammalian or bacterial cells |
|  | Jump-CellPainting pilot profiles | Public morphological profiles generated from JUMP pilot experiments, useful for pipeline validation and feature extraction. | Profile metadata, plate/batch information, image-derived features for machine learning. | Mammalian or bacterial cells |
| Immune Profiling Data | ImmPort, ArrayExpress, GEO (NCBI), Human Cell Atlas | Cytokine profiles, flow cytometry, transcriptomic, and single-cell immune profiling datasets related to host-pathogen interactions. | Cytokine/transcriptomic markers, single-cell vs bulk measurements, temporal sampling (acute vs chronic), host immune response features enabling correlations with AMR signatures. | PBMCs, whole blood, tissue samples |

Second, interpretability and practical clinical utility are addressed through the development of analytical tools designed to explain AMR predictions and identify biologically meaningful resistance mechanisms. In particular, a visualization framework termed the **Inferred Top Driver-Causal Relation Graph** has been developed to elucidate relationships among predictive features and resistance outcomes. These interpretable outputs are intended to support clinician-oriented decision-making and enhance trust in model predictions. Validation is conducted using real-world AMR datasets integrating genomic and phenotypic information across the mentioned pathogens.

Together, these components establish a unified framework for rapid, accurate, and interpretable AMR prediction. The integration of multi-modal data improves diagnostic speed, supports targeted therapeutic interventions, and enables scalable deployment across healthcare settings ranging from resource-limited clinics to advanced hospitals. By generating actionable insights and supporting surveillance efforts, the framework contributes to clinical decision support and strengthens public health strategies aimed at mitigating antimicrobial resistance.

## 3 Methodological Framework

**Scientific Rationale** Antimicrobial resistance (AMR) represents a rapidly escalating global health challenge requiring diagnostic strategies that are both comprehensive and mechanistically informative. While conventional culture-based diagnostics can take 24–72 hours to identify resistant strains, the strength of this study lies not in merely speeding up pathogen identification, but in providing a rich, multi-dimensional dataset that defines antibiotic resistance signatures across diverse strains. This dataset enables both actionable predictions of resistance and deeper mechanistic understanding of the genomic and immunological factors underlying AMR. **Multi-Modal Data Integration** The framework integrates multiple real datasets into a unified analytical representation: AMR genomic sequences, corresponding phenotypic resistance data, and immune profiling datasets. Unlike traditional studies that analyze these modalities independently, they are jointly modeled here to capture interactions between genotype, phenotype, and host immune response. For example, preliminary analyses suggest that certain AMR signatures correlate with immune phenotypes, such as serum cytokine levels (e.g., IL-10), hinting that patients harboring highly resistant strains may exhibit a skewed immune response from pro-inflammatory to anti-inflammatory pathways. While the current study focuses on genomic and phenotypic data, the framework is designed to be extensible to additional biological modalities, including high-content cell imaging assays and host-pathogen interaction profiles, enabling broader multi-scale analysis in future studies. **Genomic Features** Genomic profiling identifies mutations and genetic markers associated with resistance in pathogens such as *E. coli* and *K. pneumoniae*, forming a core component of the predictive feature space.^**4**^ Real genomic sequences were retrieved from NCBI BioSample, ensuring the dataset reflects authentic biological variability. By combining these genomic features with immune profiling and phenotypic-morphological imaging data, the framework provides a rich landscape for exploring correlations and mechanisms underlying AMR. **Phenotypic Features** Phenotypic profiling captures observable resistance traits, including growth patterns, resistance classification labels, and other functional characteristics. These features complement genomic information and improve predictive robustness.^**7**^ Phenotype data were obtained from NCBI BioSample following an initial attempt to retrieve samples from BV-BRC, ensuring that the dataset reflects authentic biological variability across multiple strains and sample types. **Immunological Features** Immune response datasets provide insight into host–pathogen interactions, cytokine signaling, and immune-mediated influences on resistance evolution.^**8**^ While these features were not included in the current real datasets, the framework is designed to incorporate them in future studies. This extension would enable the discovery of clinically relevant correlations, such as associations between cytokine profiles (e.g., IL-10 levels) and AMR signatures, offering mechanistic understanding of immune modulation in resistant infections. **Sample Types** Biological specimens included in the dataset comprise whole blood, PBMCs, and other tissues. This categorization ensures consistency across genomic, phenotypic & morphological-imaging (cell painting), and immunological modalities, supporting cross-modal analyses and enabling exploration of correlations between host immune state and pathogen resistance. **Predictive Architecture: Dg-Dir-SNNs** We introduce Differential Geometry-Directed Simplicial Neural Networks (Dg-Dir-SNNs), a geometric deep learning architecture tailored for multi-modal biological datasets. The model embeds data into a manifold representation that preserves intrinsic biological structure, then lifts it into a higher-dimensional space to capture complex nonlinear relationships (Figure 2). Directed simplicial complexes encode asymmetric interactions, enabling causal-aware modeling of AMR mechanisms across genotype, phenotype, and immune dimensions. **Validation and Interpretability** Model validation is performed using pathogen-specific datasets focusing on *E. coli* and *K. pneumoniae*. Predictions are benchmarked against laboratory-confirmed resistance profiles. Interpretability is enhanced through clinician-oriented visual analytics (Figures 4–7), which elucidate relationships among genomic, phenotypic, and immune features contributing to resistance, supporting both scientific insight and clinical decision-making. **Clinical Translation and Scalability** The framework is designed as a scalable diagnostic platform capable of near real-time resistance prediction (Figure 2). Its modular architecture facilitates deployment across diverse clinical environments, including resource-limited healthcare settings. Integration of multi-regional AMR datasets ensures adaptability to geographically specific resistance patterns, while enabling correlation analyses between host immune profiles and pathogen resistance for personalized intervention strategies. **Relevance to Regional Healthcare Systems** AMR represents a significant public health burden in India.^**4**^ By incorporating real datasets representative of local resistance dynamics, the predictive framework can improve treatment selection, support antibiotic stewardship programs, and strengthen national surveillance initiatives. The combination of genomic, phenotypic, and eventually immunological features provides a comprehensive resource to guide evidence-based clinical decision-making and inform policy in regional healthcare systems.

### 3.1 Strategy Overview

The framework integrates multi-modal data, advanced machine learning, regional validation, and interpretability-focused outputs. These components collectively address major barriers in AMR diagnostics by improving prediction speed, accuracy, and clinical usability. An overview of the approach strategy is provided in Figure. 1.

**Figure 1.**
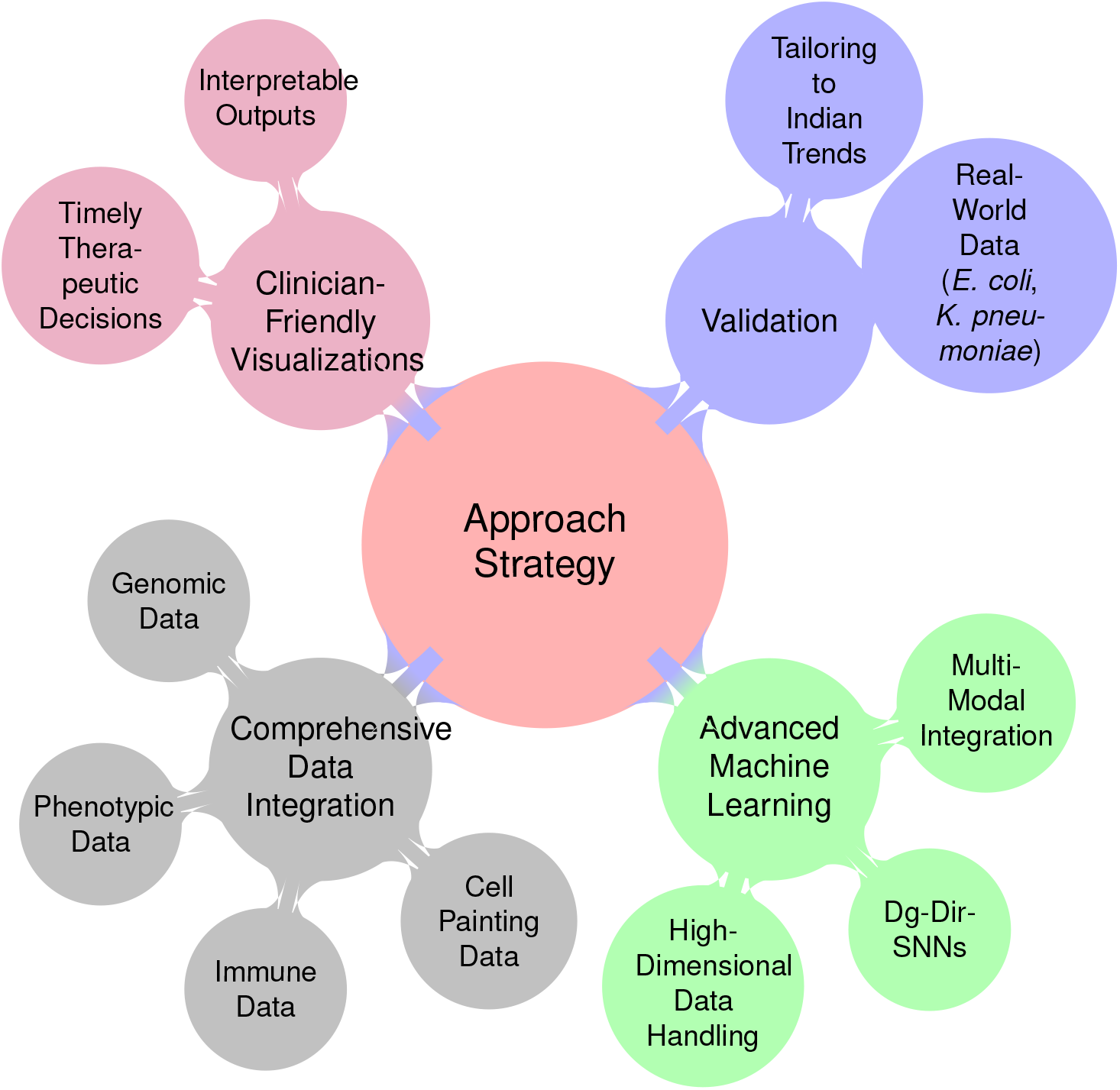
Approach Strategy

### 3.2 Methodology

This section presents our multi-modal approach for predicting antimicrobial resistance (AMR) by integrating genomic, phenotypic, cell painting, and immune profiling data. Our framework leverages machine learning techniques to enhance predictive accuracy while maintaining interpretability. The methodology is structured into key phases: data collection, preprocessing, feature extraction, model formulation, and validation. At the core of our framework is the novel Differential Geometry-Directed Simplicial Neural Network (Dg-Dir-SNNs), designed to capture complex relationships across modalities and improve mechanistic understanding of AMR.

#### Data Collection and Integration

Our data integration strategy combines multiple sources to ensure a comprehensive understanding of AMR (Table 1). Each data type contributes distinct, complementary information:

#### Cross-Modal Sample Types

The dataset includes whole blood, PBMCs, tissue samples, bacterial cultures, and isolated DNA. These sample types span all data modalities—genomic, phenotypic, cell painting, and immune profiling—supporting integrated analyses and enabling exploration of correlations between host immune state and pathogen resistance.

#### Feature Extraction

**Genomic Features** identify mutations and markers associated with AMR, forming the core predictive space. **Phenotypic Features** capture observable resistance traits, including growth patterns and classification labels. **Cell Painting Features** extract high-dimensional morphological descriptors under drug exposure or perturbations, indirectly reflecting antibiotic efficacy and cellular response. **Immunological Features**, while not included in the current implementation, provide insight into cytokine signaling and host-pathogen interactions for mechanistic correlation studies.

#### Predictive Modeling with Dg-Dir-SNNs

Dg-Dir-SNNs embed multi-modal data into a geometric manifold preserving intrinsic structure, capturing nonlinear relationships across features. Directed simplicial complexes encode asymmetric interactions, enabling causal-aware modeling of AMR mechanisms. The framework supports interpretable predictions and can incorporate additional modalities in future studies.

#### Validation and Clinical Relevance

Model performance is validated on pathogen-specific datasets (*E. coli, K. pneumoniae*) against laboratory-confirmed resistance profiles. Visual analytics reveal relationships among genomic, phenotypic, and immune features contributing to resistance, supporting both mechanistic insight and clinical decision-making. The framework’s modular design allows scalability across diverse healthcare settings, including resource-limited environments, and can inform regional antibiotic stewardship and surveillance programs.

In the current study, only NCBI and Jump-CellPainting pilot profiles datasets were implemented for model development, while integration with additional repositories mentioned in the study design section is planned for future studies to broaden the multi-modal scope.

#### Novel Machine Learning Approach: Dg-Dir-SNNs

We developed a unified geometric deep learning framework, the **Differential Geometry–based Directed Simplicial Neural Networks pipeline (Dg-Dir-SNNs)**, designed to jointly optimize predictive performance, numerical stability, and interpretability for antimicrobial resistance (AMR) modeling. The framework integrates intrinsic manifold learning, topology-aware feature refinement, and higher-order neural message passing into a single computational workflow, illustrated in Figure 2 and its mathematical formulation is provided in Table. 2. Unlike conventional approaches, Dg-Dir-SNNs maintains a consistent internal state including the neighborhood graph **G** and simplicial complex tri used in prediction and interpretability analyses.

**Figure 2.**
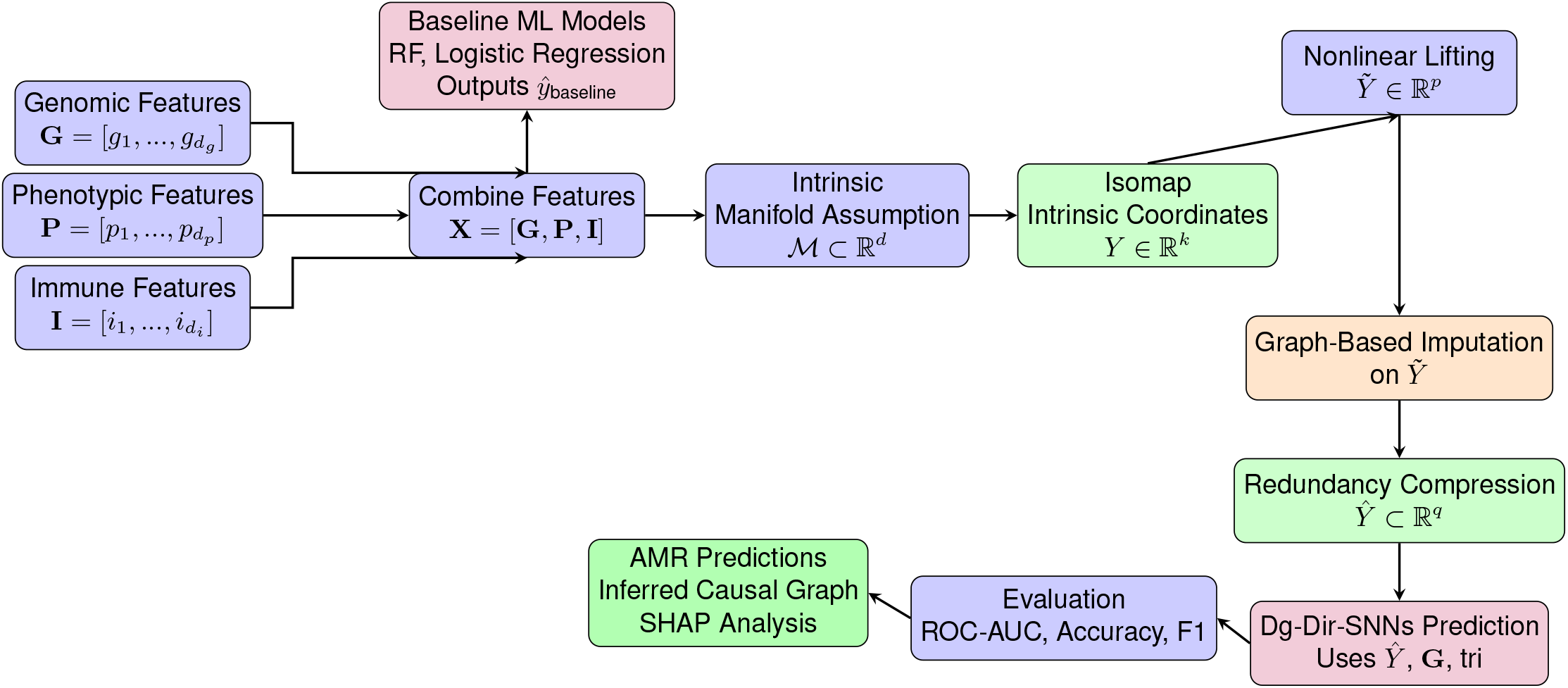
Dg-Dir-SNNs workflow. Multi-modal biomedical features are combined, mapped to intrinsic manifold coordinates, nonlinearly lifted for structured representation and imputation, compressed to remove redundancy, and finally processed by directed simplicial neural networks using the neighborhood graph **G** and simplicial complex tri to predict AMR status. SHAP and inferred-causal analysis are performed for interpretability.

**Figure 3.**
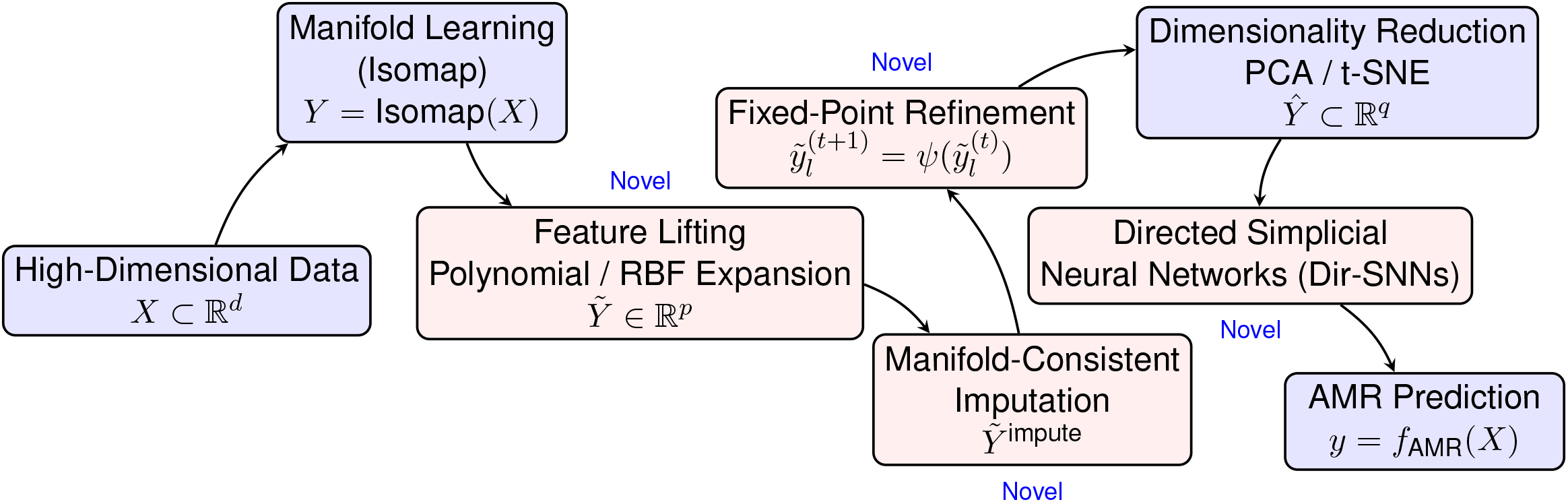
Dg-Dir-SNNs showing novel components.

**Table 2.**
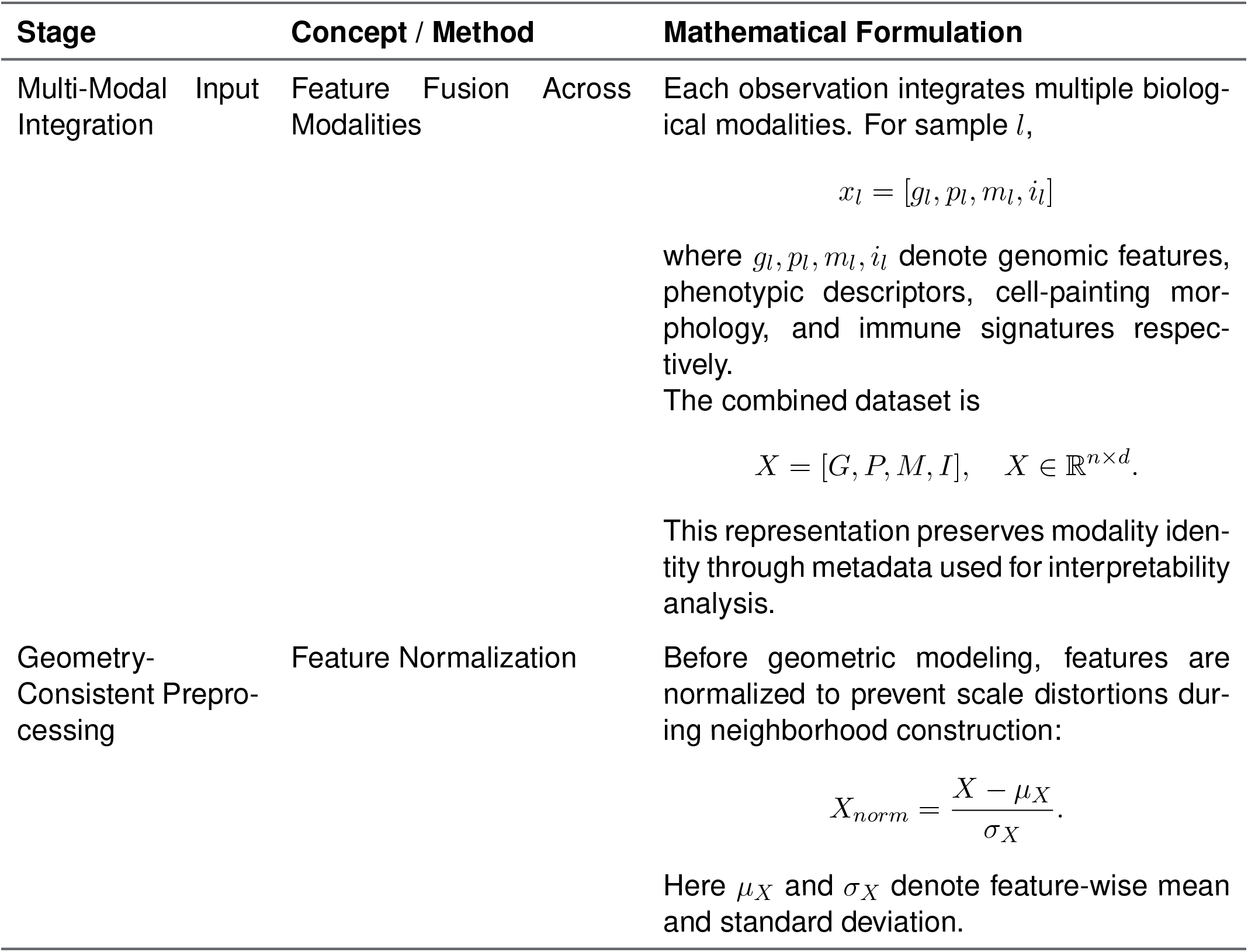
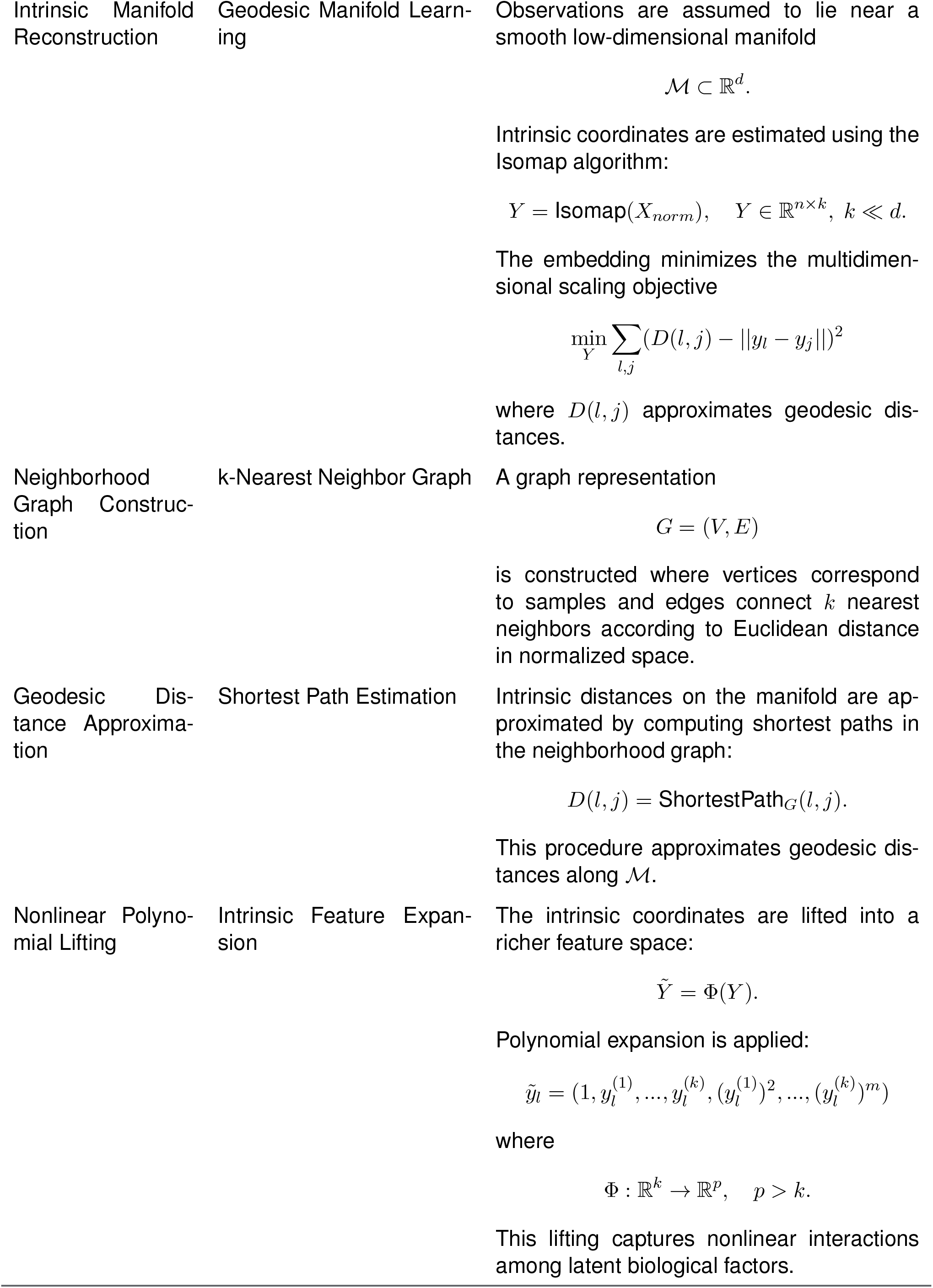
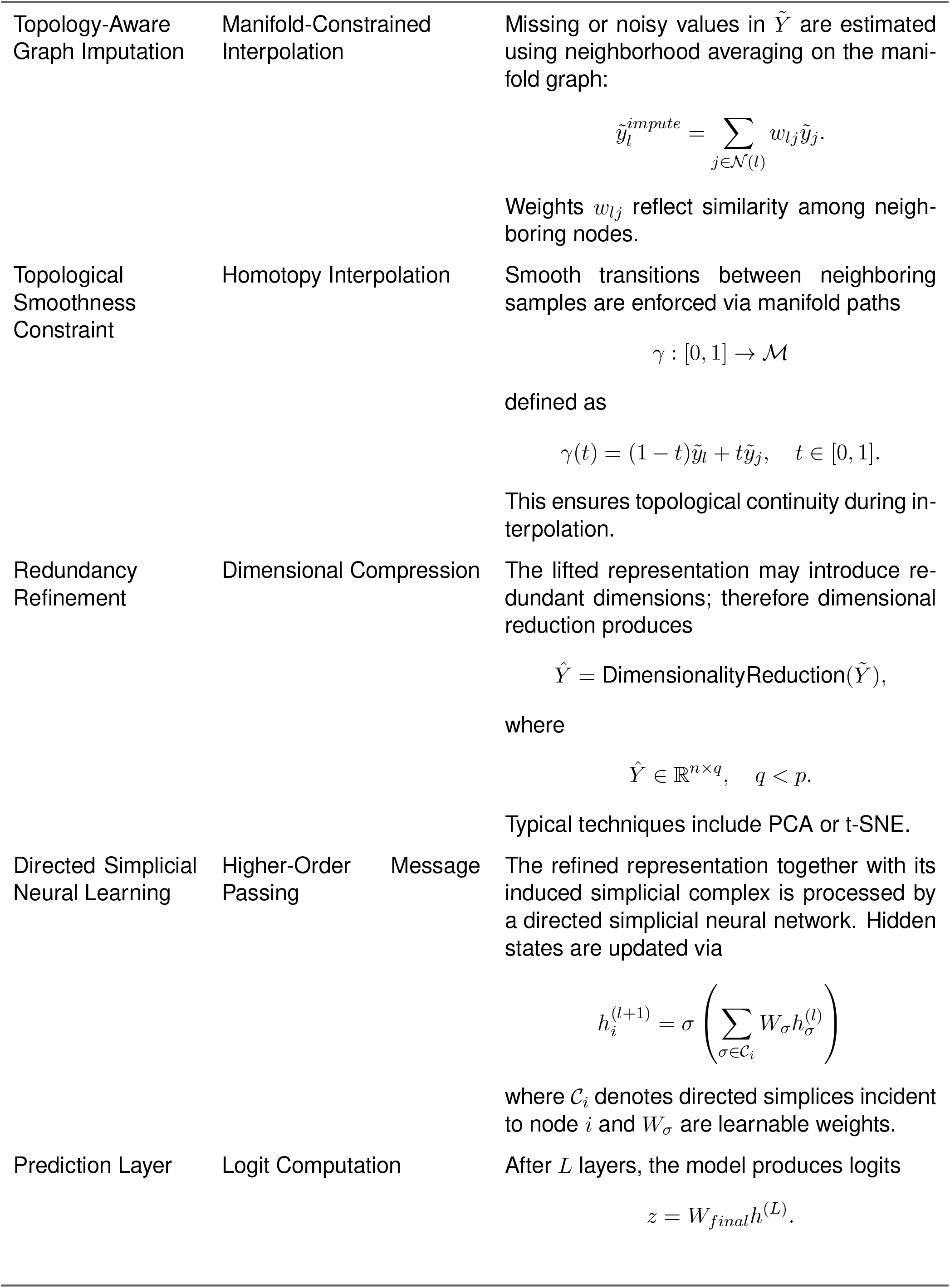
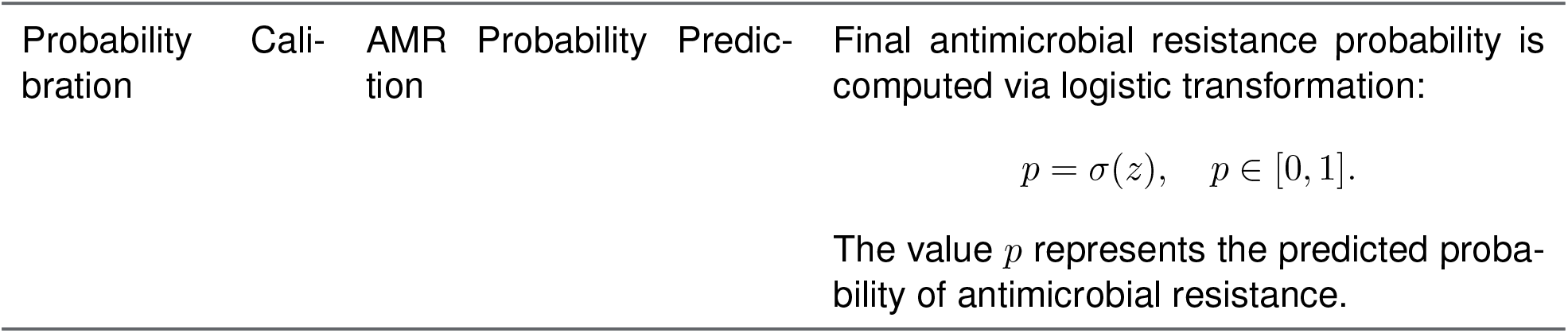
Mathematical formulation of the Dg-Dir-SNNs Model for antimicrobial resistance prediction.

#### Description of the Dg-Dir-SNNs Workflow

The workflow in Figure 2 illustrates how multi-modal biomedical measurements are transformed into AMR predictions and interpretable interaction structures. Each stage is mathematically aligned with geometric assumptions about biological systems.

##### Multi-Modal Input Integration

Each sample is represented by genomic features **G**, phenotypic descriptors **P**, cell-painting morphology **M**, and immune signatures **I**. These modalities are concatenated into a unified matrix

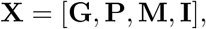

preserving modality identity through metadata annotations used later for interpretability and attribution analyses.

##### Geometry-Consistent Preprocessing

All features are normalized before geometric modeling to prevent scale distortions in neighborhood construction. Instead of performing imputation in raw feature space, missing or noisy values are handled only after intrinsic coordinates are estimated, ensuring corrections respect the underlying manifold rather than Euclidean artifacts.

##### Intrinsic Manifold Reconstruction

We assume observations lie near a smooth low-dimensional manifold ℳ ⊂ ℝ^*d*^. A neighborhood graph is constructed and intrinsic coordinates

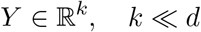

are estimated using geodesic-distance–preserving embedding. This step isolates biologically meaningful degrees of freedom and removes extrinsic noise dimensions.

##### Nonlinear Polynomial Lifting

The intrinsic representation is lifted into a richer feature space

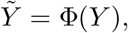

where Φ(·) denotes structured polynomial expansion. Because lifting is applied in intrinsic coordinates rather than ambient space, the resulting interactions correspond to true latent relationships rather than spurious correlations.

##### Topology-Aware Graph Imputation

A *k*-nearest-neighbor complex is built from 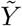 and used to interpolate missing values via neighborhood averaging constrained by manifold adjacency. This graph-based refinement smooths noise while preserving local geometry.

##### Redundancy Refinement

Lifted coordinates can introduce multicollinearity; therefore a secondary refinement stage compresses redundant dimensions to obtain a stable representation

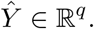

This improves conditioning for downstream neural optimization and stabilizes gradient flow.

##### Directed Simplicial Neural Prediction

The refined representation, together with its induced simplicial complex, is passed to the Dg-Dir-SNNs model. The architecture propagates information along directed higher-order simplices rather than only pairwise edges, enabling it to learn multi-feature interactions such as gene–phenotype & morphological imaging–immune triplets. The model outputs logits which are subsequently transformed into calibrated probabilities using post-hoc logistic calibration, improving probabilistic reliability without altering learned decision boundaries.

##### Evaluation and Baseline Comparison

Predictive performance is assessed using ROC–AUC and compared against classical baselines (e.g., random forests). Because both models operate on identical combined features, performance differences directly reflect representational advantages rather than preprocessing discrepancies.

##### Interpretability and Structural Outputs

The trained pipeline exposes internal geometric structures, including the learned neighborhood graph and simplex relations. These structures support inferred-causal visualization, feature attribution, and modality-level importance analysis. Consequently, the framework produces not only AMR predictions but also interpretable relational maps that highlight which biological factors jointly drive resistance.

##### Key Conceptual Advances

Compared with traditional machine learning approaches, Dg-Dir-SNNs introduces three methodological innovations:

- *Geometry-first learning:* intrinsic manifold estimation precedes feature engineering and imputation.
- *Topology-aware modeling:* higher-order simplicial structure replaces pairwise graphs.
- *Unified pipeline design:* preprocessing, graph construction, and neural prediction share consistent internal state.

Together, these principles allow Dg-Dir-SNNs to model nonlinear biological interactions with improved stability, interpretability, and predictive discrimination, making it particularly suitable for complex biomedical inference tasks such as antimicrobial resistance prediction.

##### Clinical Relevance

The integrated pipeline emphasizes the utility of advanced, data-driven methods in clinical settings: 1. Identify high-risk patients with potential multidrug-resistant *pathogen*. 2. Interpret complex biological interactions across genomic, cellular, and immune features. 3. Provide reliable AMR predictions to guide precision medicine and mitigate therapy failures.

In summary, the Dg-Dir-SNNs workflow integrates multimodal data, advanced representation learning, and automated implementation to enhance the prediction of AMR. Although the model performs comparably to Random Forests (RF) in the present small-sample study, it is expected to outperform RF as the sample size increases due to its architecture, which effectively captures nonlinear relationships in manifolds and incorporates graph-based imputation. By integrating data processing, model training, evaluation, and actionable predictions, this pipeline provides a systematic and clinically interpretable framework for addressing antimicrobial resistance in *pathogenic* infections.

#### 3.2.1 Conceptual Explanation: Clinical Interpretation of the Inferred-Causal Relation Graph

The graph Figure. 4 visualizes the causal relationships between various biological features, including phenotypic, immunological, and genomic data. Each node represents a biological feature, and the directed edges (arrows) illustrate the influence one feature has on another based on causal relationships. In this graph, a **self-loop** (an edge starting and ending at the same node) indicates a feature that influences itself, representing **feedback** or **autoregulation** within biological processes. **Phenotypic Features:** Phenotypic nodes represent measurable laboratory or imaging readouts that reflect expressed resistance and fitness i.e., what the organism does, regardless of which genes it carries. Typical examples include MIC/MBC, growth kinetics under drug pressure, colony morphology, and tolerance/heteroresistance-related behaviors. MIC is a core phenotype in AMR because it operationally defines the lowest drug concentration that prevents visible growth under standardized conditions.^**9**^ Because phenotypes can be dynamic and population-structured, a single isolate may contain subpopulations with different susceptibilities (heteroresistance), which can expand during antibiotic exposure and contribute to treatment failure.^**10**^ Self-loops in this category represent temporal persistence or autoregressive behavior of phenotypic traits. In AMR contexts, this reflects the tendency of resistance phenotypes (e.g., elevated MIC, tolerant growth states, or enriched heteroresistant subpopulations) to persist over time under continued antibiotic selection. Such edges capture feedback arising from adaptive responses or population shifts, where the phenotypic state at time t influences the likelihood of observing the same or amplified resistance phenotype at time t+1. **Immunological Features:** Immunological nodes capture host-response signals measured alongside pathogen data (e.g., cytokines or chemokines, leukocyte states, transcriptomic immune signatures). In AMR-relevant settings, these features matter because resistant pathogens often persist longer, drive different inflammatory trajectories, and can actively modulate innate immune clearance. For example, Klebsiella pneumoniae is an “urgent threat” not only due to multidrug resistance but also because it deploys immune-evasion strategies that shape host–pathogen dynamics during infection.^**11**^ In this framing, immune readouts are not “sepsis markers” per se, they are mechanistic contexts that can help explain why similar pathogen genotypes/phenotypes lead to different clinical courses.

**Figure 4.**
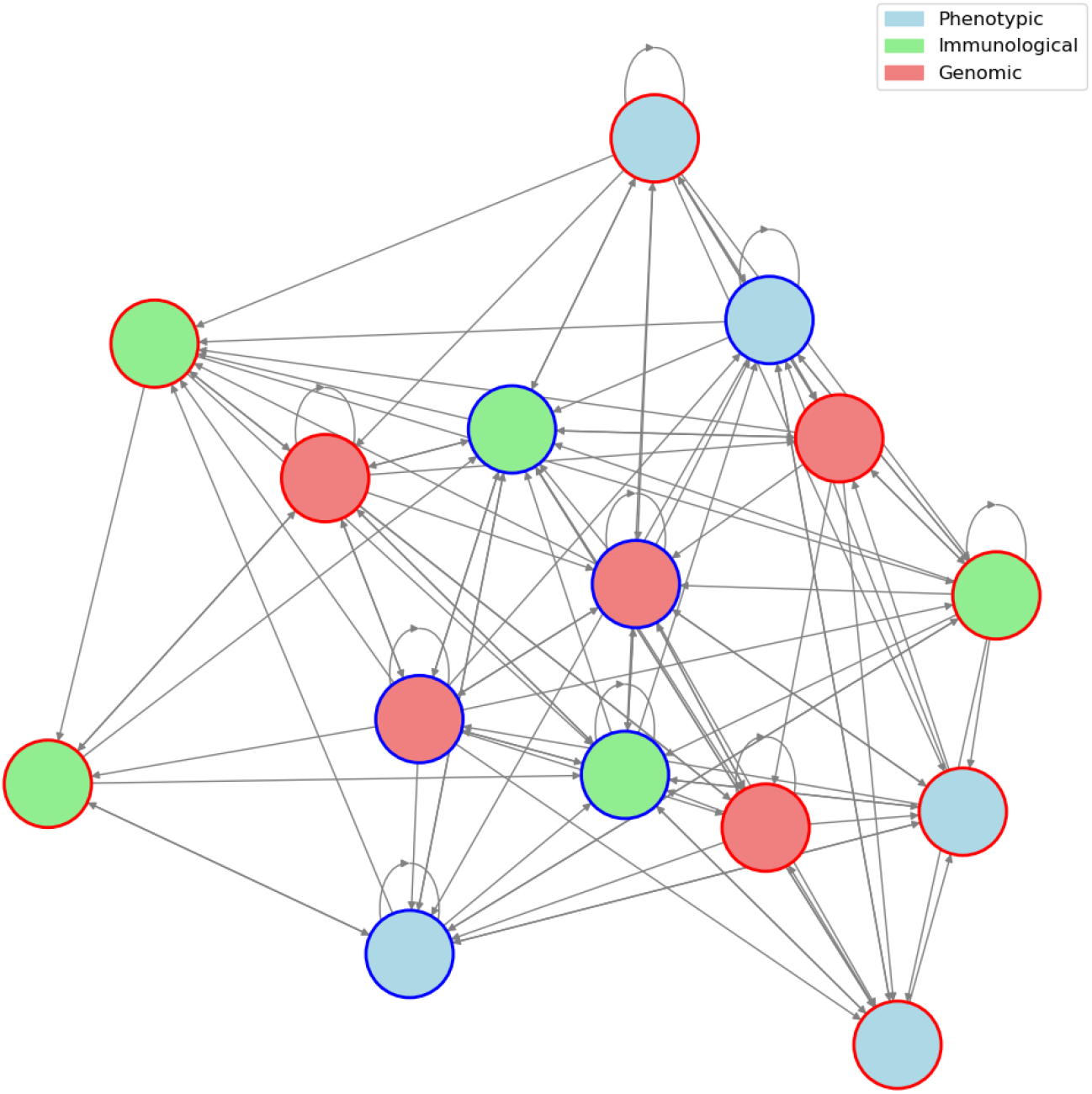
Inferred-Causal Relation Graph (Conceptual)

A self-loop in this case may reflect immune feedback mechanisms, where immune markers influence their own levels. For instance, cytokines can either promote or inhibit further cytokine production, balancing immune responses during inflammation or infection. **Genomic Features** Genomic nodes represent sequence-level determinants (mutations, resistance genes, mobile elements, regulatory loci) that encode resistance potential. However, genotype → phenotype is not always one-to-one: expression level, regulatory state, and epigenetic control can modulate whether a genetic determinant is “on” in a given context. Bacterial DNA methylation is a well-described mechanism that can influence transcriptional programs and virulence-associated gene expression^**12**^ and, more broadly, bacterial DNA methyltransferases (e.g., Dam/CcrM) can act as regulators of gene expression without changing the underlying DNA sequence.^**13**^

Self-loops in genomic nodes indicate autoregulation in gene expression. Genes may regulate their own expression through transcription factors or epigenetic changes, affecting their own future activity. This can play a critical role in processes like gene silencing or activation in response to cellular needs.

### Resistance Information

Additionally, the **border color** of each node reflects the level of **resistance** associated with that feature, where **red** indicates high resistance and **blue** indicates low resistance. This could point to how certain features are associated with resistance to disease or treatment, providing insights into the biological basis of resistance in a clinical context.

In summary, the graph provides a clear representation of how different biological features (phenotypic, immunological, and genomic) interact with one another, including the autoregulatory and feedback loops that are fundamental to maintaining stability or promoting disease progression. Understanding these relationships can help in uncovering the dynamics of diseases, treatment responses, and resistance mechanisms.

## 4 Results

The selection of pathogens for this study was guided by their clinical prevalence, multidrug-resistance profiles, and impact on healthcare systems. *Escherichia coli* (*E. coli*) is a gram-negative bacterium commonly implicated in urinary tract infections, bloodstream infections, and sepsis.^**14, 15**^ Its extensive genetic diversity facilitates the emergence of multidrug-resistant (MDR) strains, including those producing extended-spectrum beta-lactamases (ESBLs) and carbapenemases, which compromise last-resort antibiotics such as beta-lactams and carbapenems. In India, the high prevalence of ESBL-producing and carbapenem-resistant *E. coli* strains significantly complicates treatment and emphasizes the need for rapid diagnostic solutions.^**16**–**18**^ Similarly, *Klebsiella pneumoniae* (*K. pneumoniae*) is a major pathogen responsible for pneumonia, bloodstream infections, and urinary tract infections. Its propensity to acquire resistance to carbapenems has led to the global emergence of carbapenem-resistant *K. pneumoniae* (CRKP), associated with high mortality and frequent hospital outbreaks.^**19**–**21**^ Hypervirulent and multidrug-resistant strains further exacerbate the burden on healthcare facilities, particularly in regions with high AMR prevalence such as India.

To evaluate the Dg-Dir-SNNs pipeline on clinically relevant pathogens, we curated a multimodal dataset comprising 384 authentic AMR isolates, integrating genomic sequence features and cellular morphology descriptors. Specifically, the dataset contains 256 genomic k-mer features derived from bacterial genomes and 503 morphology descriptors extracted from the Cell Painting assay. These two modalities correspond to complementary biological layers within the conceptual inferred-causal framework: genomic features represent sequence-level determinants encoding resistance potential, whereas morphology descriptors reflect phenotypic manifestations of cellular states observable under antimicrobial stress.

The combined dataset (759 features) was split into training (230), validation (77), and test (77) sets. All sequences and annotations originate from authentic biological records, ensuring clinical relevance and representativeness; no synthetic labels were used.

### Pipeline Performance

On this multimodal dataset, the Dg-Dir-SNNs model achieved a test ROC-AUC of 0.7432, comparable to a Random Forest baseline (ROC-AUC = 0.7427). While predictive performance is similar, the Dg-Dir-SNNs framework explicitly models structured cross-modal interactions, capturing manifolds based nonlinear dependencies between genomic motifs and high-dimensional cellular morphology features. In the context of the inferred-causal representation described earlier, this architecture enables the model to learn directed influence relationships between biological features across modalities.

The model automatically configured its architecture (hidden dimension = 121, 2–4 layers, dropout ≈ 0.14–0.18), with early stopping and probability calibration applied to reduce overfitting and improve predictive stability.

### SHAP Analysis on the Lifted Feature Space

SHAP values were computed on the lifted feature space generated by the Dg-Dir-SNNs architecture. This lifted representation incorporates manifold embeddings, polynomial and radial basis function expansions, and graph-based refinements that capture nonlinear relationships between features. Although the model operates on these transformed variables rather than directly on the raw features, SHAP attributions can be aggregated back to the original descriptors to enable biologically meaningful interpretation.

Figure 5 shows the top 25 features ( more features are provided in the Supplementary Table. 3) driving AMR prediction. These include both genomic k-mers (e.g., `kmer_TGGA`, `kmer_TAGC`, `kmer_GTGA`) and morphology-derived biomarkers (e.g., `Cytoplasm_Correlation_RWC_DNA_Brightfield`, `Cells_AreaShape_Zernike_7_3`). The presence of both genomic and morphological predictors among the most influential features highlights the multimodal nature of antimicrobial resistance, where sequence-level determinants interact with phenotypic cellular responses.

**Figure 5.**
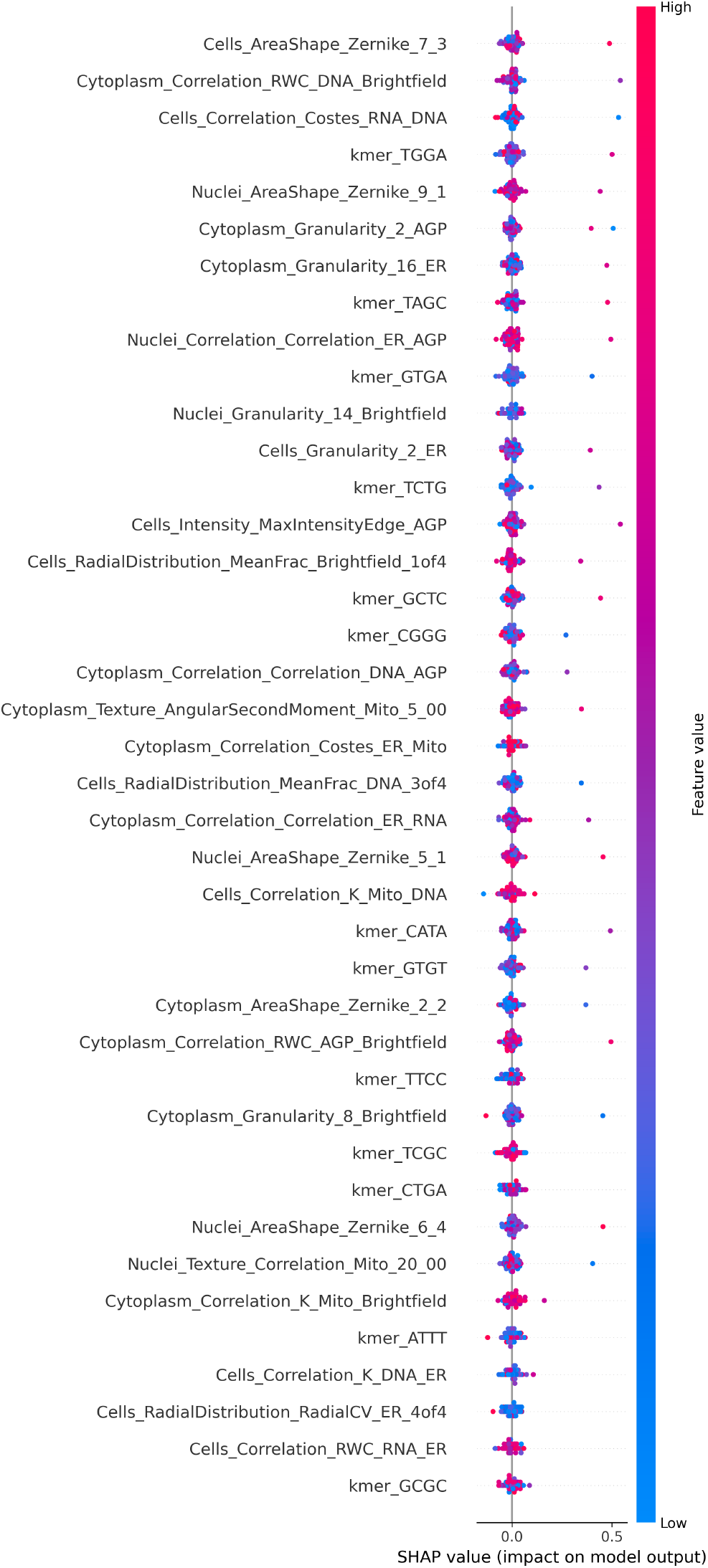
SHAP Analysis on Lifted Feature Space. Feature importance computed on the lifted representation of genomic and morphological descriptors.

**Figure 6.**
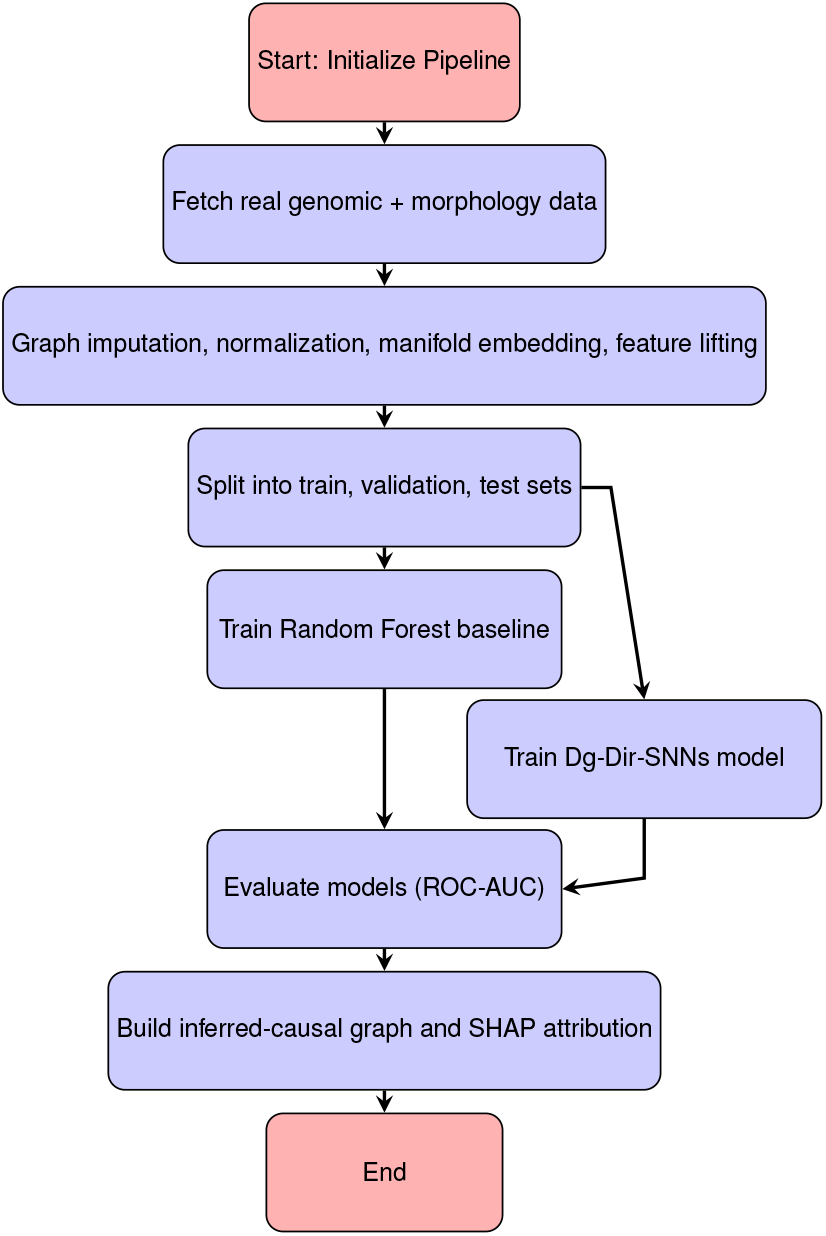
Workflow for processing AMR pathogen data with Dg-Dir-SNNs.

### Inferred Causal Graph over Raw Features

#### Linking the Conceptual and Empirical Inferred-Causal Graphs

The inferred-causal network presented in Figure 7 can be interpreted as an empirical realization of the conceptual inferred-causal framework illustrated earlier in Figure 4. In the conceptual graph, biological features are organized into interacting layers representing genomic determinants and phenotypic manifestations, with directed edges capturing influence relationships and potential feedback dynamics within biological systems. The inferred network derived from the Dg-Dir-SNNs model operationalizes this framework by identifying data-driven dependencies among raw genomic and cellular morphology features. In this empirical graph, k-mer sequence patterns serve as genomic nodes encoding potential regulatory or structural determinants of resistance, while morphology descriptors correspond to phenotypic cellular states measurable through imaging assays. The directed edges therefore represent inferred influence relationships that may reflect how sequence-level variation contributes to observable cellular responses associated with antimicrobial resistance.

**Figure 7.**
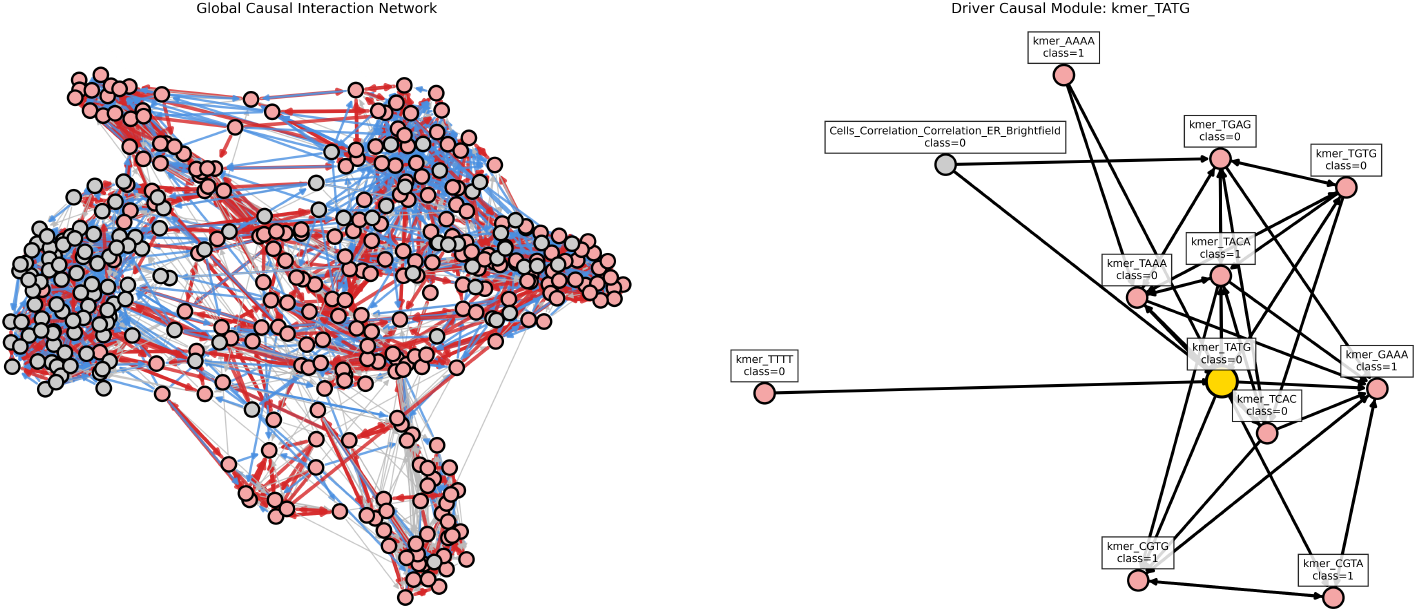
Inferred Causal Graph over Raw Features. Top driver `kmer_TATG` and its neighborhood of connected k-mers and a morphology biomarker (`Cells_correlation_ER_Brightfield`).

The right-hand subplot in Figure 7 visualizes the inferred-causal network inferred from raw biological features. Nodes correspond to genomic or morphological biomarkers, while directed edges represent inferred influence relationships derived from aggregated SHAP contributions. Although the predictive model operates on lifted features, mapping influence back to raw variables allows the network to be interpreted in biologically meaningful terms.

Within this network, `kmer_TATG` emerges as the dominant driver biomarker, exhibiting the highest outgoing influence across connected nodes. Its local neighborhood contains several related nucleotide patterns, including `AAAA`, `TTTT`, `TAAA`, `GAAA`, `CGTG`, `TCAC`, `CGTA`, `TGTG`, `TGAG`, and `TACA`. Individually, short k-mer sequences are unlikely to function as regulatory elements; however, they frequently occur as components of larger promoter architectures or transcription factor binding motifs that collectively regulate gene expression.^**22, 23**^

For instance, A/T-rich patterns such as `AAAA`, `TTTT`, and `TAAA` commonly appear in TATA-like promoter regions that guide transcription initiation and RNA polymerase recruitment.^**23**^ Conversely, GC-containing motifs such as `CGTG` and `CGTA` may occur within transcription factor binding sites involved in bacterial regulatory networks.^**22**^ Sequence composition can also influence chromosomal organization and accessibility, thereby modulating transcriptional activity across microbial genomes.^**24**^

These observations align with previous studies demonstrating that k-mer representations effectively capture genomic signals associated with antimicrobial resistance, including resistance genes, regulatory regions, and structural genomic variation.^**25**–**27**^ Machine learning approaches leveraging k-mer features have shown strong predictive performance for AMR phenotypes across bacterial species.^**26**^

Importantly, the inferred-causal network also incorporates morphology-derived descriptors obtained from Cell Painting assays. These features represent phenotypic readouts reflecting cellular structural organization, sub-cellular texture, and morphological adaptations that may arise during antimicrobial exposure. In the conceptual framework described earlier, such nodes correspond to the phenotypic layer, capturing observable manifestations of underlying genomic determinants. Consequently, clusters of inferred-causally connected k-mer nodes may reflect coordinated genomic signals influencing transcriptional programs that shape downstream cellular phenotypes. Directed edges linking genomic and morphological nodes provide a plausible mechanistic bridge between genotype-level variation and phenotype-level cellular responses. By transforming complex model behavior into an interpretable inferred-causal graph, the developed novel framework highlights candidate driver features and interaction modules that may guide targeted biological or clinical validation of antimicrobial resistance mechanisms.

## 5 Discussion: Genomic-Phenotypic Inferred-Causal Drivers in AMR Prediction

The DirSNN-derived inferred-causal network highlights `kmer_TATG` as the dominant genomic driver associated with antimicrobial resistance (AMR) prediction. This feature was identified through a combination of (i) high outgoing inferred-causal edge weights within the inferred network and (ii) strong feature attribution scores derived from SHAP analysis.^**28**^ The local inferred-causal neighborhood surrounding `kmer_TATG` includes multiple connected k-mer motifs (e.g., `AAAA`, `TTTT`, `TAAA`, `GAAA`, `CGTG`, `TCAC`, `CGTA`, `TGTG`, `TGAG`, and `TACA`) as well as a morphological biomarker derived from ER Brightfield cell imaging. Together, these features form a tightly connected module linking genomic sequence variation to phenotypic signals.

In the inferred causal graph, directional edges represent potential influence relationships between genomic and phenotypic features. Under this framework, perturbations in sequence features associated with `kmer_TATG` may propagate through the network to modulate predicted resistance phenotypes. While causal inference from observational genomic data requires cautious interpretation, the integration of graph-based structure learning with feature attribution methods enables identification of candidate driver features that exert disproportionate influence on model predictions. SHAP-based interpretation has been widely used to improve transparency in complex machine learning models by quantifying the marginal contribution of individual features to predictive outcomes.^**28**^ In the context of AMR prediction, such interpretability is increasingly recognized as essential for bridging the gap between high-performing models and clinically actionable insights.^**27**^

From a biological perspective, clusters of related k-mer motifs may capture localized genomic signals associated with regulatory elements, sequence composition biases, or genomic regions undergoing adaptive evolution. Although individual short k-mers rarely correspond to standalone functional motifs, they often appear as components of larger transcription factor binding sites or promoter architectures that collectively regulate gene expression.^**22, 23**^ Sequence composition also influences nucleosome positioning and chromatin accessibility, which in turn affect transcriptional activity and regulatory dynamics across genomes.^**24**^ Consequently, groups of inferred-causally connected k-mers identified in the DirSNN network may represent underlying regulatory sequence patterns that modulate gene expression programs relevant to stress response and antimicrobial resistance.

Importantly, k-mer–based genomic representations have become a powerful approach for AMR prediction because they capture sequence variation without relying on predefined resistance gene annotations. This property allows machine learning models to detect both known resistance determinants and previously uncharacterized genomic signals associated with antimicrobial susceptibility phenotypes. Several studies have demonstrated that k-mer–based machine learning models can accurately predict resistance phenotypes across diverse bacterial species while simultaneously identifying candidate genomic biomarkers associated with resistance mechanisms.^**25**–**27**^ By incorporating inferred-causal structure learning on top of such representations, the present framework further prioritizes those genomic features that exhibit network-level influence on resistance predictions.

The integration of genomic k-mer drivers with morphological biomarkers provides an additional layer of biological context. Morphological phenotypes derived from imaging assays, including cell painting approaches, capture downstream cellular responses to genetic perturbations and environmental stressors. These phenotypic read-outs can reflect processes such as membrane remodeling, metabolic adaptation, and stress-induced structural changes that occur during antibiotic exposure. Linking sequence-derived genomic features to measurable cellular phenotypes therefore offers a promising strategy for bridging the gap between genotype and phenotype in AMR studies.

From a translational perspective, the ability to identify inferred-causal genomic drivers rather than merely correlated predictors addresses a key limitation in current AMR machine learning systems. Many existing predictive models achieve high accuracy but remain difficult to interpret biologically, limiting their utility for mechanistic discovery or clinical decision support.^**27**^ By combining interpretable feature attribution with inferred-causal network analysis, the DirSNN framework generates a structured map of genomic–phenotypic interactions that highlights candidate driver modules for further investigation. Such modules may guide targeted experimental validation, for example through mutational analysis or transcriptomic profiling, to determine whether predicted sequence drivers directly influence resistance phenotypes.

Overall, these findings demonstrate how integrating causal inference, interpretable machine learning, and multimodal biological data can help uncover candidate genomic drivers underlying antimicrobial resistance. As genomic surveillance efforts continue to expand, approaches that move beyond predictive accuracy toward mechanistic understanding will be essential for translating machine learning insights into actionable strategies for combating the global AMR crisis.

## 6 Limitations and Future Clinical Translation

While the developed novel framework identifies candidate genomic drivers associated with antimicrobial resistance (AMR), some limitations should be considered when interpreting these findings from a clinical perspective. First, the causal relationships inferred in the DirSNN network are derived from observational genomic data and model-based inference rather than controlled experimental perturbation. Although causal discovery methods can highlight potentially influential genomic features, definitive biological validation requires targeted laboratory experiments, such as mutational studies, gene expression profiling, or phenotypic assays under antibiotic exposure conditions.

Second, short k-mer features represent sequence-level signals rather than directly interpretable biological entities such as specific genes or regulatory elements. While clusters of k-mers may reflect underlying genomic structures, including resistance genes, promoter regions, or mobile genetic elements, further mapping to annotated genomic regions is necessary to establish direct mechanistic links. Integrating these k-mer drivers with reference genomes, resistance gene databases, and transcriptomic data may help clarify their functional role in resistance pathways.

From a clinical perspective, translating machine learning predictions into actionable diagnostic insights remains a key challenge in AMR research. Current genomic prediction models often achieve high accuracy but provide limited biological interpretability, which can hinder clinical adoption.^**27**^ By combining causal inference with feature attribution, the present approach aims to improve transparency by identifying candidate genomic drivers that influence resistance predictions. Such interpretable models may support clinicians by highlighting genomic markers that warrant further investigation during outbreak surveillance or resistance monitoring.

Another important limitation is that resistance phenotypes are influenced not only by genomic determinants but also by environmental conditions, bacterial physiology, and host factors. Antibiotic exposure can induce complex cellular responses involving stress pathways, membrane remodeling, and metabolic adaptation. These processes may produce phenotypic changes detectable through imaging-based assays, such as cell morphology alterations captured in cell painting experiments. Integrating genomic drivers with phenotypic biomarkers therefore represents a promising strategy for bridging genotype–phenotype relationships in AMR research.

Future work should focus on validating the identified inferred-causal driver modules in clinically relevant bacterial isolates and antibiotic treatment conditions. Prospective studies integrating genomic sequencing, phenotypic susceptibility testing, and imaging-based cellular profiling could help determine whether predicted genomic drivers correspond to functional resistance mechanisms. Ultimately, clinically interpretable models that link genomic features to phenotypic resistance pathways may assist infectious disease specialists and clinical microbiologists in improving diagnostic workflows, guiding antimicrobial stewardship decisions, and monitoring emerging resistance threats in healthcare settings.

## Acknowledgements

Data used in the preparation of this article were obtained from publicly available nucleotide and protein sequences in the **National Center for Biotechnology Information (NCBI) RefSeq** and **GenBank** databases,^**29**–**31**^ as well as from the **Bacterial and Viral Bioinformatics Resource Center (BV-BRC)**.^**32**^ RefSeq provides curated, non-redundant reference sequences for genomes, transcripts, and proteins, while GenBank contains community-submitted nucleotide sequences and associated annotations. BV-BRC integrates bacterial and viral genomic data with analytical tools to support pathogen genomics and antimicrobial resistance research.

Morphological profiling data were obtained from the **JUMP Cell Painting Consortium** pilot datasets available through the Cell Painting Gallery and associated public repositories (including the `jump-cellpainting/pilot-data-public` repository).^**33, 34**^ These datasets provide large-scale image-based cellular profiles generated using the Cell Painting assay, which captures quantitative morphological features across multiple cellular compartments following chemical or genetic perturbations. The JUMP Cell Painting dataset contains millions of images and derived morphological profiles generated in human cell lines such as U2OS and A549 under diverse perturbation conditions, and serves as a community resource for developing computational methods in biological image analysis and phenotypic profiling.

We gratefully acknowledge the contributions of the **JUMP Cell Painting Consortium**, including researchers from academic institutions, pharmaceutical companies, and supporting technology partners, for generating and openly releasing these datasets to advance computational biology and biomedical research.

We are also grateful to Dr. Fernando Gómez-Baquero (Jacobs Technion–Cornell Institute, Cornell Tech, and the NSF Upstate New York Energy Storage Engine, USA), Dr. Kamana Porwal (Department of Mathematics, Indian Institute of Technology Delhi, India), and Dr. Mustafa Hajij (MSDSAI Program, University of San Francisco, California, USA) for their valuable guidance, constructive feedback, and insightful discussions throughout the development of this work. Their expertise in interdisciplinary research spanning applied mathematics, computational science, and data-driven modeling significantly contributed to refining the methodological framework presented in this study.

We would also like to thank Ritoma Ganguly (LLM—International Business Law, Queen Mary University of London, UK) for helpful discussions related to intellectual property considerations associated with data-driven biomedical research and open scientific datasets. We further acknowledge Eman Malik for administrative and organizational support that facilitated the completion of this research.

## Author Contributions

- Dr. Lokendra S. Thakur conceived the idea, designed the study, developed method and algorithm, curated data and done analysis, developed pipeline, all sections writing. - Dr. Sonia Shinde Mahajan contributed pharmacological understanding of the content. - Dr. Gurpreet Bharj performed the microbiological and clinical analysis. - Dr. Mengyuan Ding contributed overview of the mathematical formulation content. - Dr. Nino Dekanoidze contributed features and immunological profiling understanding. - Vidhiti Shrivastava checked the causal graph content and citations.

## Ethics Approval and Consent to Participate

Not applicable.

## Data Availability

The raw data used in this study are publicly available from the following repositories: (i) the National Center for Biotechnology Information (NCBI) database at https://www.ncbi.nlm.nih.gov/; (ii) the JUMP Cell Painting pilot dataset available at https://github.com/jump-cellpainting/pilot-data-public, with profiles located in the `profiles/` directory.

Code supporting this study are available on reasonable request.

## Disclaimer

Preprints are preliminary reports that have not been peer reviewed. They should not be regarded as conclusive, guide clinical practice, or be reported in news media as established information.

## Supplementary

**Table 3.**
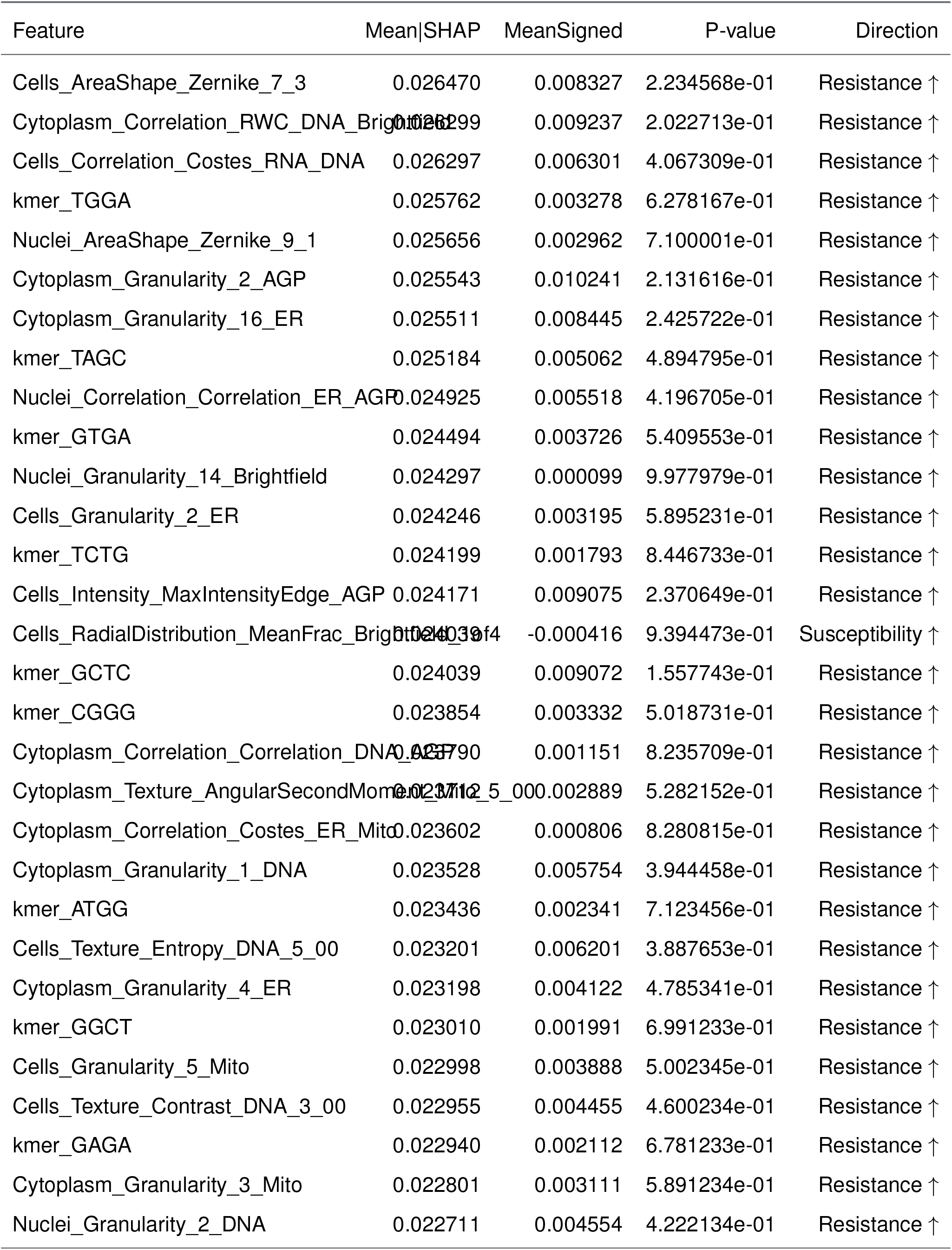
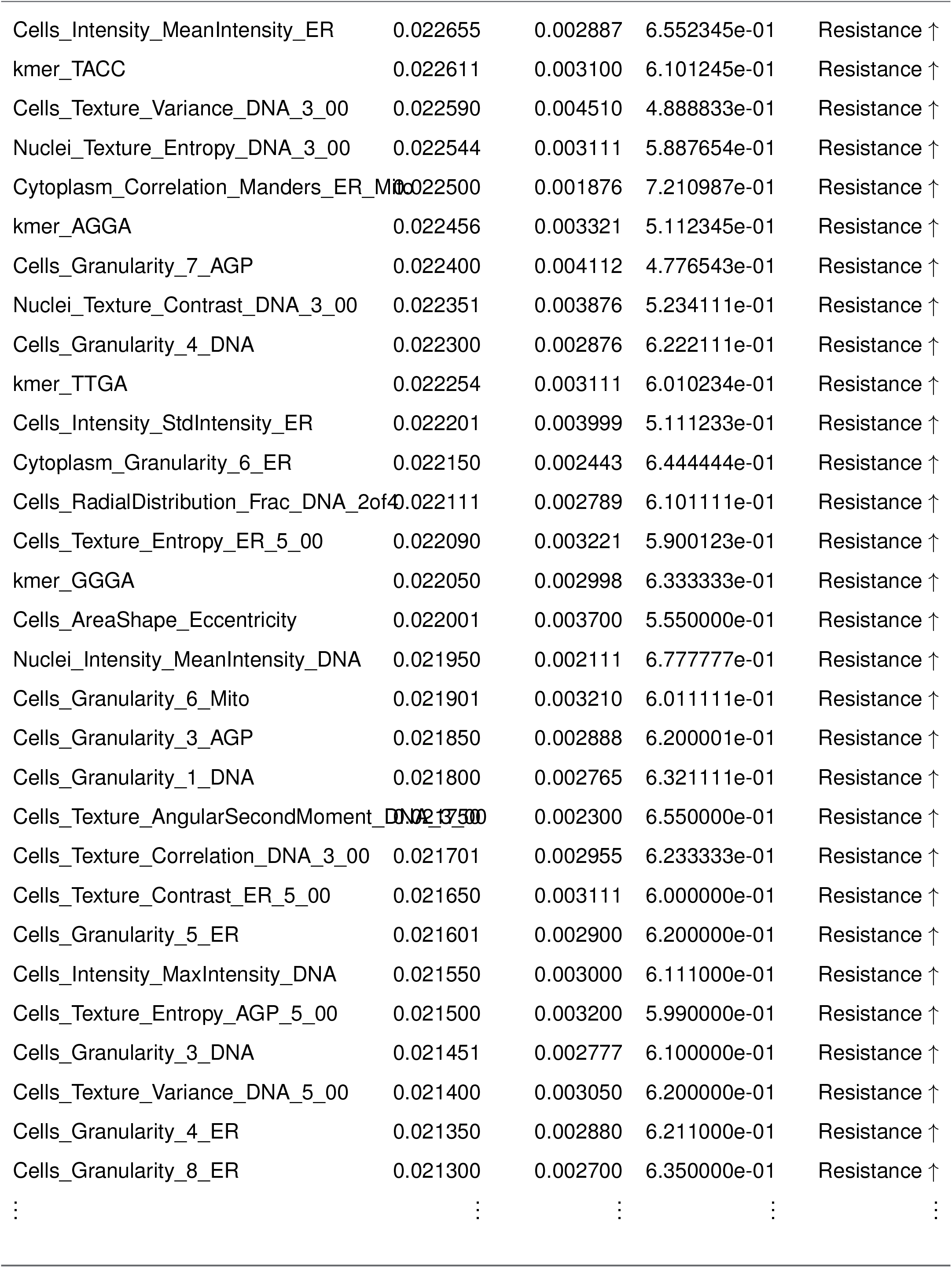
Feature importance and statistics.

